# A multi-modal transcriptomic atlas reveals the cellular and spatial landscape of canine gastric cancer

**DOI:** 10.64898/2026.06.19.732976

**Authors:** Shawna R. Cook, Jessica Z. Schneider, Rebecca M. Harman, Elaine A. Ostrander, Paul J. J. Mandigers, Jacquelyn M. Evans

## Abstract

Gastric cancer is the fifth leading cause of cancer-related mortality in humans globally and remains a clinical challenge with limited treatment options and poor survival. Dogs develop spontaneous gastric cancer that parallels the clinical presentation and histology of human disease, supporting their value as a comparative oncology model. Here we present a comprehensive transcriptomic characterization of canine gastric cancer through single-nucleus RNA-sequencing, bulk RNA-sequencing, and Visium HD 3′ spatial transcriptomics of treatment-naive tumor and normal stomach tissues from Belgian Tervuren and Belgian Sheepdogs. Across 107,085 nuclei, we identified 44 distinct cell populations, including tumor-enriched states as well as profound depletion of the normal parietal and chief cell gastric lineages. Cell-cell communication analysis revealed enhanced epithelial-fibroblast crosstalk driving epithelial-mesenchymal transition. Bulk RNA-sequencing further identified enrichment of signaling pathways implicated in *H. pylori* associated human gastric carcinogenesis, including Hippo, PI3K-Akt, and Wnt. Notably, we observed cell-type-specific altered expression of *KLHL29*, *PDZRN3*, and *PLAU*, which are among our previously identified canine gastric cancer susceptibility genes, linking germline risk to specific tumor cellular contexts. These data establish the first transcriptomic atlas of canine gastric cancer and demonstrate substantial molecular homology between canine and human disease.

## INTRODUCTION

Gastric cancer is a major global health burden, ranking as the fifth leading cause of human cancer-related deaths worldwide, with an overall 5-year survival rate of 20-40%^1,2^. Naturally occurring gastric cancer in dogs is clinically and histologically similar to its human counterpart, with poor prognosis and limited treatment options^3,4^. Clinical signs, such as vomiting, hyporexia, and weight loss, typically present at advanced disease stages^4^. There remains an urgent need to identify biomarkers and effective therapies in both humans and dogs, and the parallel disease process in canines may serve as a valuable model for understanding gastric cancer biology and advancing human health.

We previously identified genetic risk factors in the predisposed Belgian Tervuren and Belgian Sheepdog breeds, which are on average 45 times more likely to be diagnosed with gastric cancer compared to other breeds^5–7^. Our genomic analyses identified 18 loci influencing susceptibility in these breeds, including well-known human gastric cancer genes such as *PTEN* and the major histocompatibility complex, as well as new associations with *PDZRN3* and *KLHL29*^8^. This work established the value of spontaneous canine gastric cancer as a genetic model for revealing novel risk genes for investigation in the human disease counterpart. However, the transcriptomic landscape of gastric cancer has not yet been explored in dogs.

In humans, single-cell RNA-sequencing has illuminated the cellular heterogeneity and complex tumor microenvironment of gastric cancer^9–11^. Here, we expand the molecular characterization of canine gastric cancer and profile tumor gene expression at single-cell resolution, generating a single-nucleus RNA-sequencing atlas and bulk transcriptomic data for tumor and normal stomach tissues from Belgian Tervuren and Belgian Sheepdogs. We apply spatial transcriptomics to resolve tissue architecture at near single-cell resolution, providing tissue-level context for cell populations. Together, these data highlight the transcriptomic homology between canine and human gastric cancer and nominate diagnostic and therapeutic candidates for further investigation.

## RESULTS

### A single-nucleus transcriptomic atlas of canine gastric cancer and normal stomach tissue

To characterize the cellular landscape of canine gastric adenocarcinoma, snRNAseq data were generated from four histologically normal stomach and seven treatment-naive primary tumor tissues collected from Belgian Tervuren and Belgian Sheepdogs. Following removal of low-quality nuclei (Methods), 107,085 nuclei were retained for analysis (66,524 tumor, 40,56 normal) with an average of 30,889 reads per nucleus and a median of 2,417 genes and 5,620 transcripts per nucleus. Data were normalized via SCTransform, integrated across samples using reciprocal principal component analysis (PCA), and clustered to identify cell populations through differential expression and canonical marker gene analysis. Unsupervised clustering identified 29 clusters representing seven broad cell type categories: epithelial, fibroblast, smooth muscle, immune, neural, endothelial, and adipocyte (Figure 1). Adipocytes were primarily observed in two tumor samples and absent from normal samples and were thus removed from downstream analyses. Compositional analysis revealed significant differences between tumor and normal samples, with depletion of epithelial (p=0.0073) and neuroendocrine (p=0.0030) cells and expansion of immune cells (p=0.0045) and fibroblasts (p=0.0058) in tumor samples (Supplemental Figure 1). Subclustering within each major cell type category resolved 44 distinct cell populations (Supplemental Figure 2), recapitulating cell types previously described in human gastric cancer single-cell studies^9–13^.

**Figure 1.**
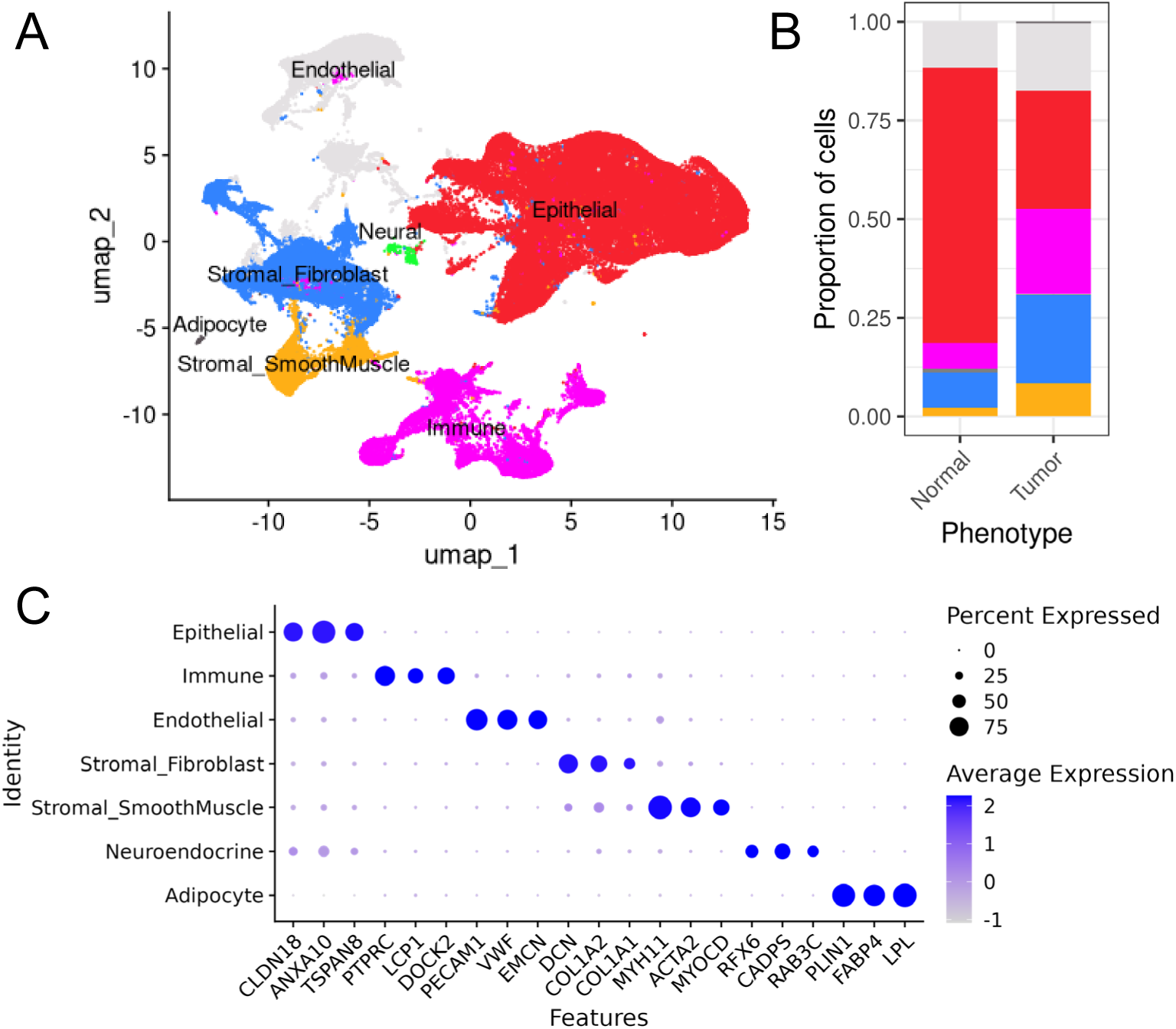
Major cell type clustering in normal (n=4) and tumor (n=7) canine stomach epithelium. (A) UMAP showing the seven broad cell populations identified in snRNAseq from 107,085 nuclei. (B) Proportion of major cell types across the normal and tumor samples. (C) Dot plot showing expression of major cell type marker genes. Dot size represents the percentage of nuclei in the cluster expressing that gene and color corresponds to average standardized gene expression.

### Epithelial cells exhibit loss of gastric identity and epithelial to mesenchymal transition signatures in tumors

Subclustering of 53,042 epithelial cells identified 11 distinct clusters capturing gastric epithelial cell type lineages (Figure 2A-C). Tumor samples exhibited dramatic depletion of differentiated gastric lineages, with loss of parietal (p_adj_ < 5×10^-5^), chief (p_adj_ < 5×10^-5^), and tuft cells (p_adj_ = 2.5×10^-4^). Concurrently, tumor samples demonstrated expansion of EMT-like (p_adj_ < 5×10^-5^), cycling (p_adj_ < 5×10^-5^), progenitor (p_adj_ = 1.25×10^-4^), and tumor-epithelial (p_adj_ < 5×10^-5^) populations (Supplemental Figure 3).

**Figure 2.**
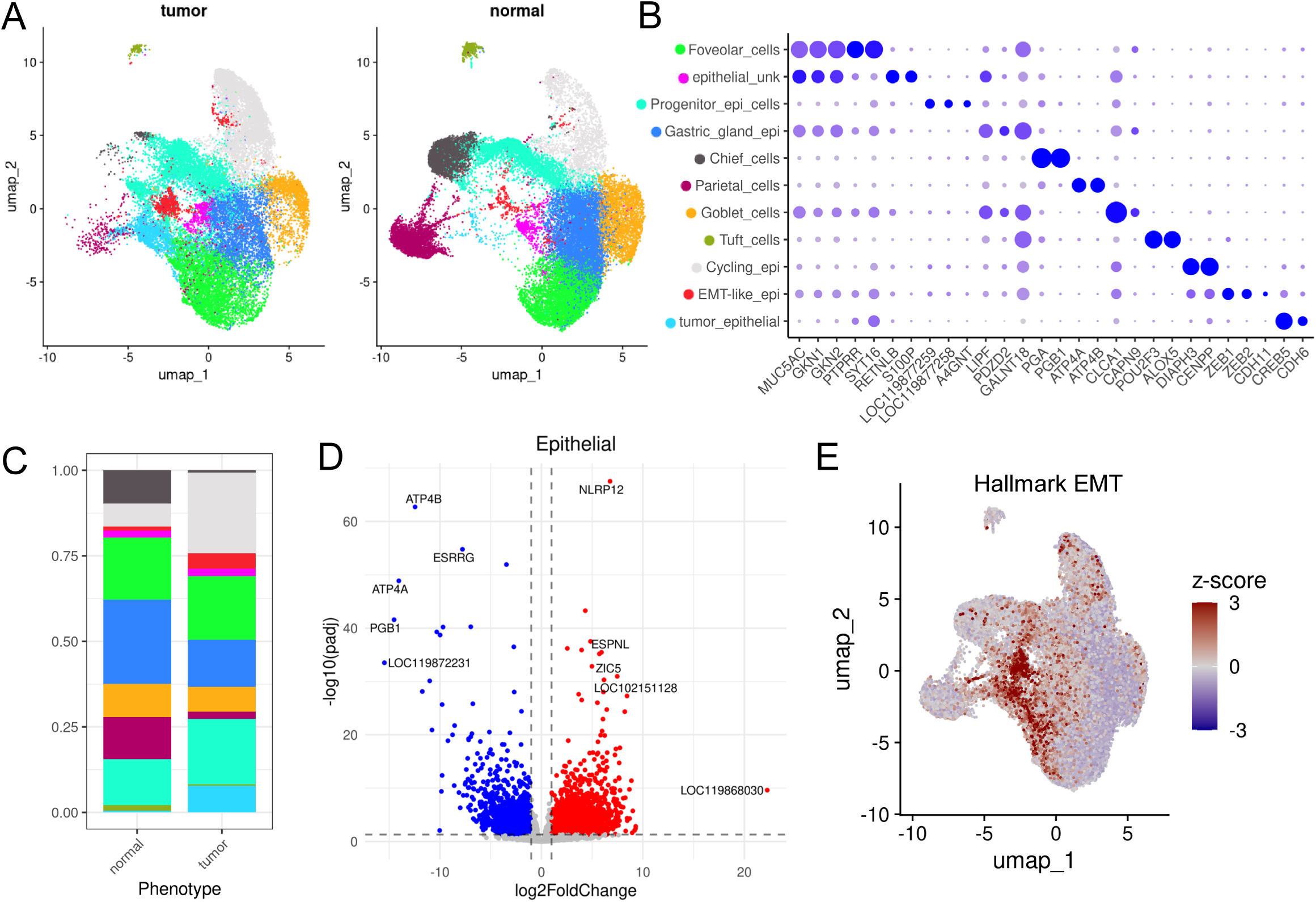
Epithelial cell type clustering and differential expression. (A) UMAP, split by phenotype, showing the 11 epithelial cell types identified across 53,042 nuclei. (B) Dot plot showing expression of epithelial marker genes. (C) Proportion of epithelial cell types across the normal and tumor samples. (D) Volcano plot illustrating differentially expressed genes from pseudobulked epithelial data. Positive Log2FoldChange indicates genes upregulated in tumor (red), while negative values (blue) represent downregulation in the tumor. (E) UMAP of the epithelial cells showing co-expression of all genes in the Hallmark epithelial-mesenchymal transition pathway. High z-score (red) indicates upregulation of pathway genes.

Pseudobulk differential gene expression analysis across all epithelial cells identified 4,553 significantly dysregulated genes (p_adj_<0.05 and |log2FC| ≥ 1), including 2,882 upregulated and 1,671 downregulated in tumor samples (Figure 2D; Supplemental Table 1). These changes recapitulated the loss of gastric epithelial identity at the gene level, with extreme reduction of the parietal cell markers *ATP4A* (log2FC = -14.01), *ATP4B* (log2FC = -12.45), and *ESRRG* (log2FC = -7.7) in tumor samples. The chief cell pepsinogen genes *PGA* (log2FC = -8.74) and *PGB1*, or *PGBP*, (log2FC = -14.52) were similarly downregulated, consistent with the loss of chief cells in tumor samples. Among upregulated genes, *NLRP12* was the most significantly dysregulated (p_adj_ = 3.07×10^-72^; log2FC = 6.77). *CDH6* was widely expressed in gastric cancer samples, with the strongest expression in tumor-epithelial cells, and was significantly upregulated relative to normal samples in which expression was nearly absent (p_adj_ = 9.70×10^-29^; log2FC = 6.11). *CDH11* was also significantly upregulated (p_adj_ = 3.36×10^-8^; log2FC = 1.97) and predominantly expressed in EMT-like epithelial cells.

Gene set enrichment analysis revealed integrin signaling (p_adj_=2.24×10^-5^) and extracellular matrix receptor interaction (p_adj_=4.02×10^-5^) as the top enriched KEGG pathways in tumor samples (Supplemental Table 2). Analysis using the MSigDB Hallmark gene sets identified epithelial-mesenchymal transition as the top enriched pathway (p_adj_= 1.67×10^-21^), consistent with the integrin and ECM signaling enrichments (Supplemental Table 3). The EMT signature was strongest in EMT-like cells expressing canonical mesenchymal transcription factors (*ZEB1*, *ZEB2*, *RBMS3*) and in tumor-epithelial cells (Figure 2E). Tumor samples additionally showed enrichment for TNF-α signaling via NFκB (p_adj_ = 1.64×10^-11^) and reduction of oxidative phosphorylation (p_adj_ = 9.19×10^-10^). Multiple human gastric cancer gene sets were significantly enriched in the canine cohort, including signatures associated with advanced stage tumors (Supplemental Table 4)^14,15^.

### Fibroblasts promote epithelial-mesenchymal transition

Subclustering of 17,050 fibroblasts resolved seven distinct populations, including multiple cancer-associated fibroblast (CAF) subtypes (Figure 3A-C). Tumor samples had significantly more pericytes (p_adj_ = 0.013) and *C3*^+^ CAFs (p_adj_ = 0.0075) and fewer epithelial-like fibroblasts (p_adj_ = 0.0018) than normal samples (Supplemental Figure 4). A rare, specialized population of *CCL19^+^*/*CXCL13*^+^ immunofibroblasts was observed in three of seven tumor samples, comprising 1.4-2.6% of fibroblast cells in those tumors. These CAF subtypes parallel populations described in human gastric single-cell studies, including the matrix CAFs and immunofibroblasts^9,16^.

**Figure 3.**
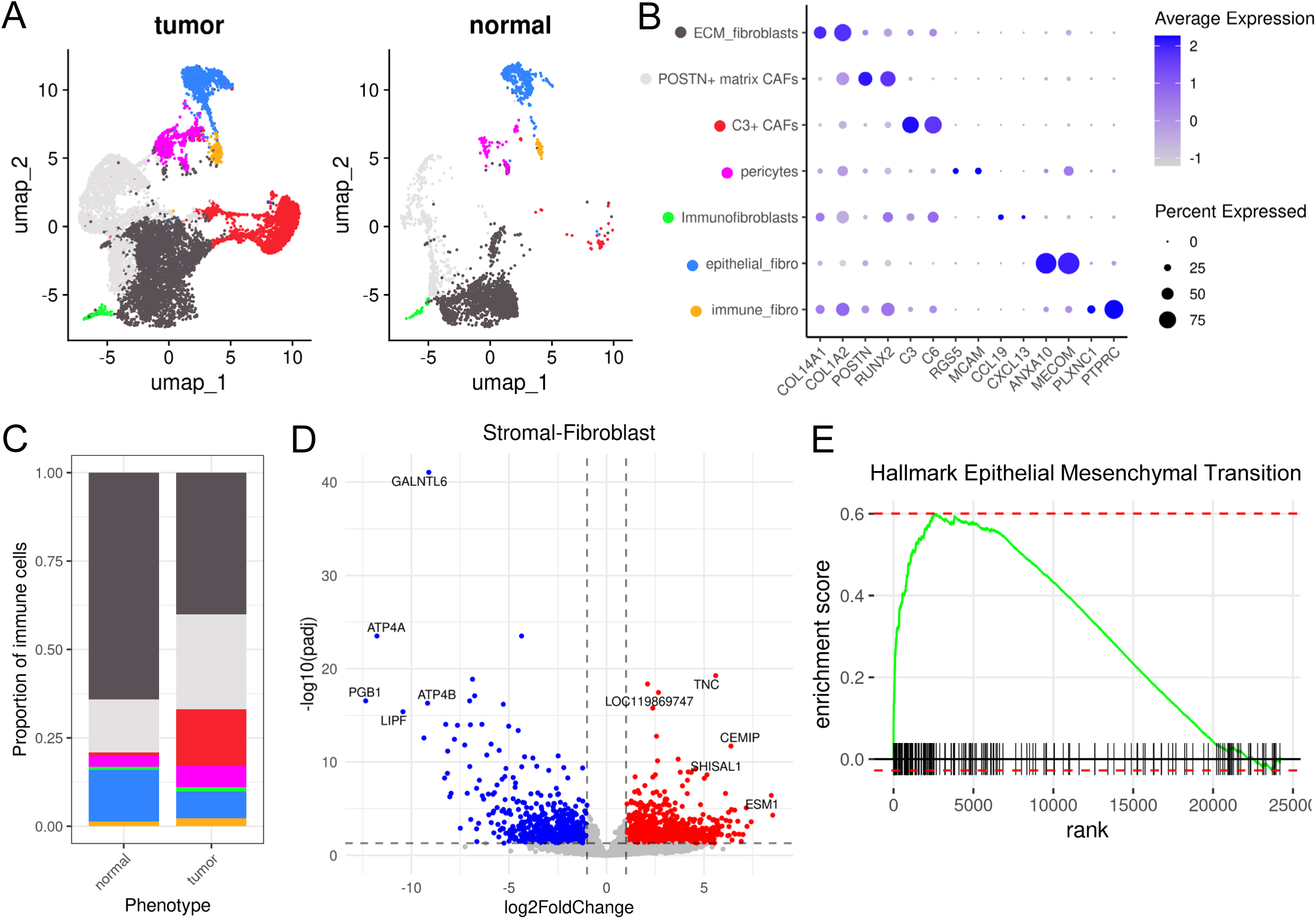
Fibroblast cell clustering and differential gene expression. (A) UMAP, split by phenotype, showing the 11 fibroblast cell types identified across 17,050 nuclei. (B) Dot plot showing expression of fibroblast marker genes. (C) Proportion of fibroblast cell types across the normal and tumor samples. (D) Volcano plot showing differentially expressed genes of the pseudobulked fibroblast nuclei data. Positive log2FoldChange (red) indicates genes upregulated in tumor and negative values (blue) represent downregulation in tumor. (E) Gene set enrichment plot of the Hallmark epithelial-mesenchymal transition pathway. Vertical black bars represent the rank of each gene in the pathway, the green line represents the running enrichment score, and the dashed lines show the maximum and minimum enrichment score.

Pseudobulk differential gene expression analysis identified *GALNTL6* (log2FC = -9.11, p_adj_ = 8.93×10^-42^) and *TNC* (log2FC = 5.60, p_adj_ = 5.42×10^-20^) as the most significantly dysregulated genes between fibroblasts from tumor and normal samples (Figure 3D; Supplemental Table 1). Gene set enrichment analysis revealed the Hallmark epithelial-mesenchymal transition pathway as the top enriched signature in tumor sample fibroblasts (p_adj_ = 4.00×10^-20^), with leading-edge genes including multiple collagens and *POSTN* (Figure 3E; Supplemental Table 3). Two cycling cell signatures, G2M checkpoint (p_adj_ = 1.66×10^-19^) and E2F targets (p_adj_ = 7.43×10^-14^) were also significantly enriched. Additional enriched pathways include TNF-α signaling via NF-κB (p_adj_ = 2.11×10^-16^) and MYC targets V1 (p_adj_ = 8.28×10^-11^), reflecting active inflammatory and proliferative programs in tumor-associated fibroblasts^17–20^.

### Immune cells display inflammatory and immunosuppressive transcriptional programs

Subclustering of 14,271 immune cells resolved 15 distinct populations spanning lymphoid and myeloid lineages (Figure 4A-C). Among the dendritic cells, a subcluster expressed *XCR1*, consistent with conventional dendritic cells (cDC1), which function in activating CD8+ T cells and natural killer cells^21^. Compositional analysis revealed an altered immune microenvironment across tumor samples, with significant expansion of tumor-associated macrophages (p_adj_ < 5×10^-5^), inflammatory macrophages (p_adj_ = 8.18×10^-3^), mature dendritic cells (p_adj_ = 0.018), and regulatory T cells (p_adj_ = 0.023). Activated T cells (p_adj_ = 6.83×10^-3^), mast cells (p_adj_ = 5.00×10^-3^), myeloid dendritic cells (p_adj_ = 0.027), and epithelial-like cells (p_adj_ = 0.013) were depleted in tumor samples (Supplemental Figure 5).

**Figure 4.**
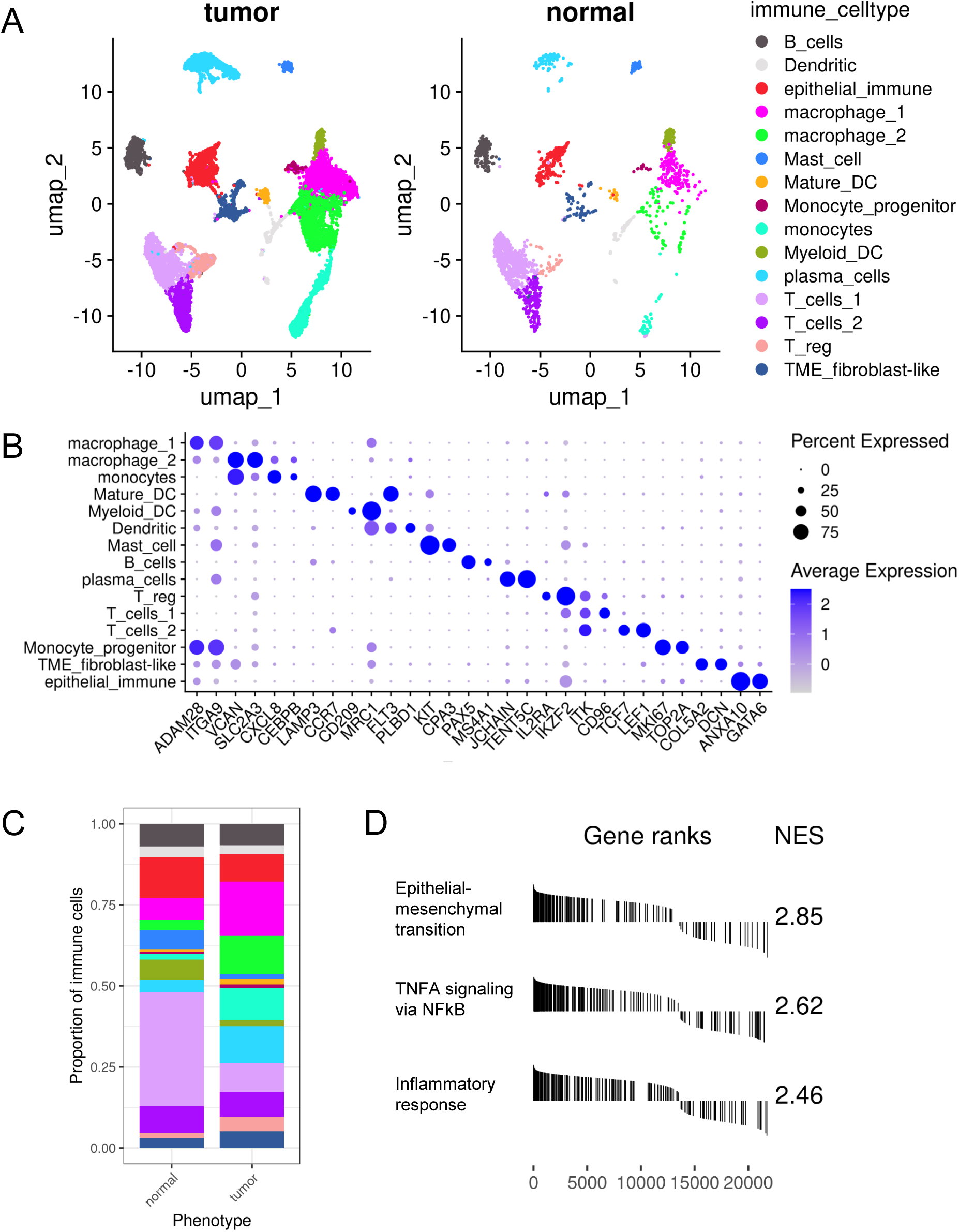
Immune cell clustering in normal and tumor canine stomach. (A) UMAP, split by phenotype, showing the 15 immune cell types identified across 14,271 nuclei. (B) Dot plot showing expression of immune marker genes. (C) Proportion of immune cell types across the normal and tumor samples. (D) Stacked plot of the top three enriched Hallmark pathways. Vertical black lines indicate the rank of each gene in its respective pathway, with those pointed upward showing upregulation and downward showing downregulation. The normalized enrichment score is shown for each pathway.

Differential gene expression analysis of the pseudobulk data revealed *CD109* (p_adj_ = 2.38×10^-11^; log2FC = 2.79), *SPP1* (p_adj_ = 1.67×10^-9^; log2FC = 4.37), and *OSMR* (p_adj_ = 5.50×10^-7^; log2FC = 2.28) as the most significantly upregulated genes in immune cells from tumor samples (Supplemental Table 1). Gene set enrichment analysis revealed strong enrichment of epithelial-mesenchymal transition (p_adj_ = 4.10×10^-28^), TNFA signaling via NFKB (p_adj_ = 3.03×10^-21^), and inflammatory response (p_adj_ = 1.40×10^-16^) pathways (Figure 4D; Supplemental Table 3). The expansion of immunosuppressive populations alongside active inflammatory signaling in tumor sample monocytes reflects the tumor-promoting inflammation paradigm characteristic of human gastric cancer^17,22^.

### Cell-cell communication analysis reveals enhanced epithelial-fibroblast crosstalk in EMT signaling pathways

Cell-cell communication analysis identified substantially increased overall intercellular signaling strength in tumor samples compared to normal. Endothelial, epithelial, and fibroblast populations exhibited the most pronounced increases in signaling strength both within and between cell types (Figure 5A-B). Among individual signaling networks, laminin, collagen, and PTPRM pathways showed the strongest tumor-specific increases (Supplemental Figure 6A). Foveolar and tumor-epithelial cells demonstrated enhanced crosstalk with both ECM-fibroblasts and *POSTN*+ matrix CAFs through the laminin (leading ligand-receptor interactions: LAMB1-CD44, LAMA2-CD44) and collagen (COL4A1/2-CD44) networks (Figure 5C). Notably, multiple genes mediating these interactions (*COL4A1*, *ITGA2*, *COL4A2*, *COL1A1*, *COL6A3*, *COL1A2*, *LAMC2*, *ITGB1*) also appeared in the leading edge of the Hallmark EMT signature, mechanistically linking epithelial-fibroblast intercellular communication to the EMT profile identified within these cell types. The PTPRM signaling architecture exhibited considerable restructuring in tumor samples, with crosstalk concentrated to and from vascular endothelial cells (Figure 5C). CAFs additionally engaged vascular endothelial cells through VEGF, with signaling driven by *VEGFA* and *VEGFC*, consistent with stromal promotion of tumor angiogenesis^9^, whereas VEGF signaling in normal cells relied on *VEGFC* and *VEGFD* (Supplemental Figure 6B).

**Figure 5.**
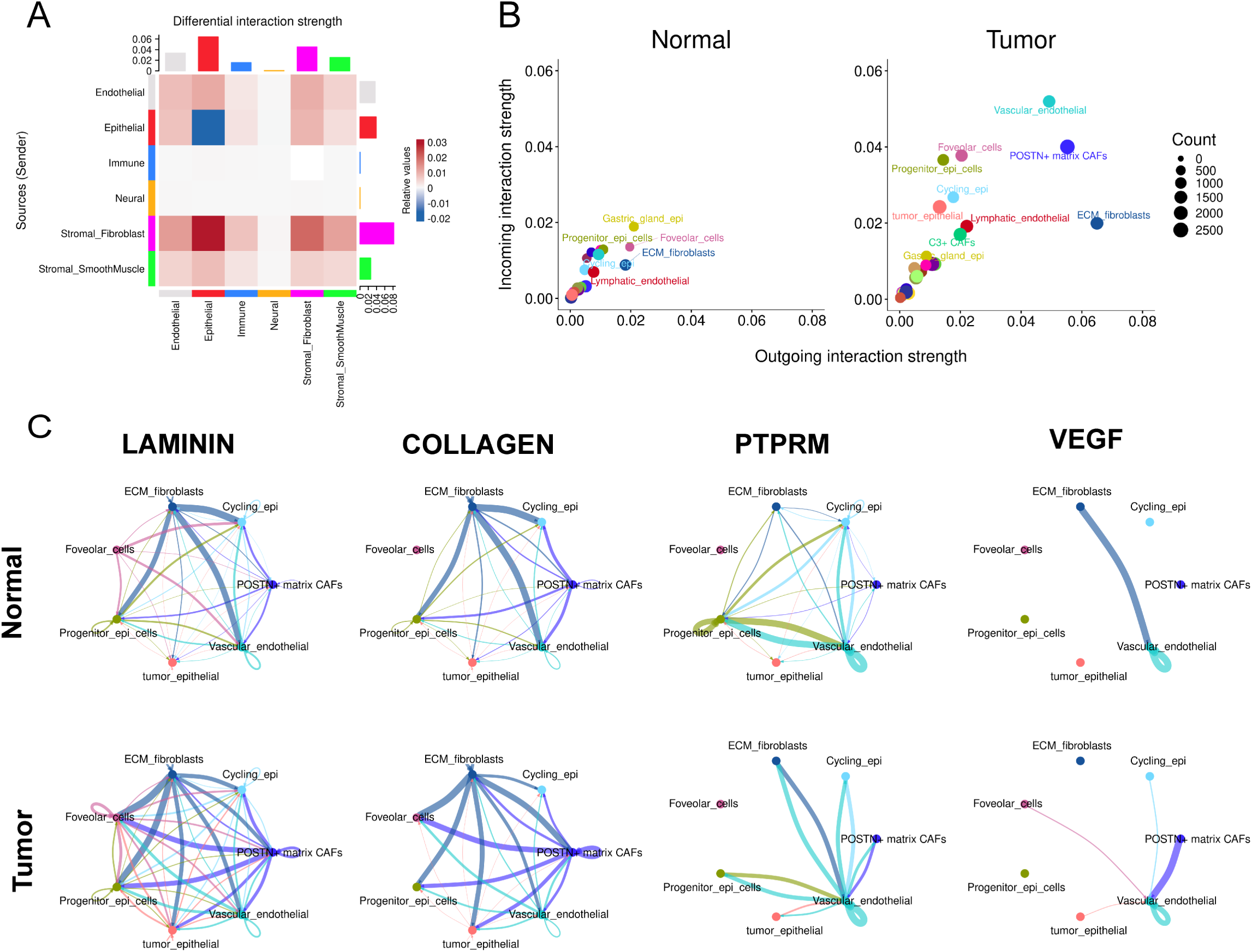
Intercellular signaling pathway analysis. (A) Heatmap illustrating differential interaction strength between broad clusters from gastric tumor and normal snRNA-seq data. Red indicates increased signaling strength in tumor as compared to normal, and blue indicates decreased signaling strength. (B) Scatter plot illustrating dominant signaling senders and receivers amongst epithelial, endothelial, and stromal fibroblast subpopulations derived from normal (left) and tumor (right) tissue. (C) Circle plots illustrating key inferred signaling networks in normal and tumor tissue. Edge color corresponds to the node from which the pathway signal originated.

### Spatial transcriptomics supports snRNA-seq findings and resolves tissue-level organization of canine gastric mucosa

Spatial transcriptomic analysis, using the Visium HD 3′ platform, of one gastric tumor and two histologically normal stomach tissues identified 15 transcriptionally distinct cell-type clusters (Figure 6A-B). Spatial organization of these clusters in normal tissue recapitulated the expected architecture of canine gastric fundic glands, with foveolar and neck cells localized toward the luminal surface and gastric gland epithelial cells comprising the basal portions of the crypts (Figure 6A), consistent with established human gastric mucosal organization^9^. Neighborhood enrichment analysis identified consistent spatial co-occurrence between surface epithelial populations (foveolar and neck cells) and between cell types residing within the gastric gland body (progenitor epithelial, parietal, and chief cells; Figure 6C). Notably, plasma cells and fibroblasts exhibited spatial co-localization with tumor-specific endothelial cells in tumor tissue, suggesting immune-stromal organization localized to the vascular niche.

**Figure 6.**
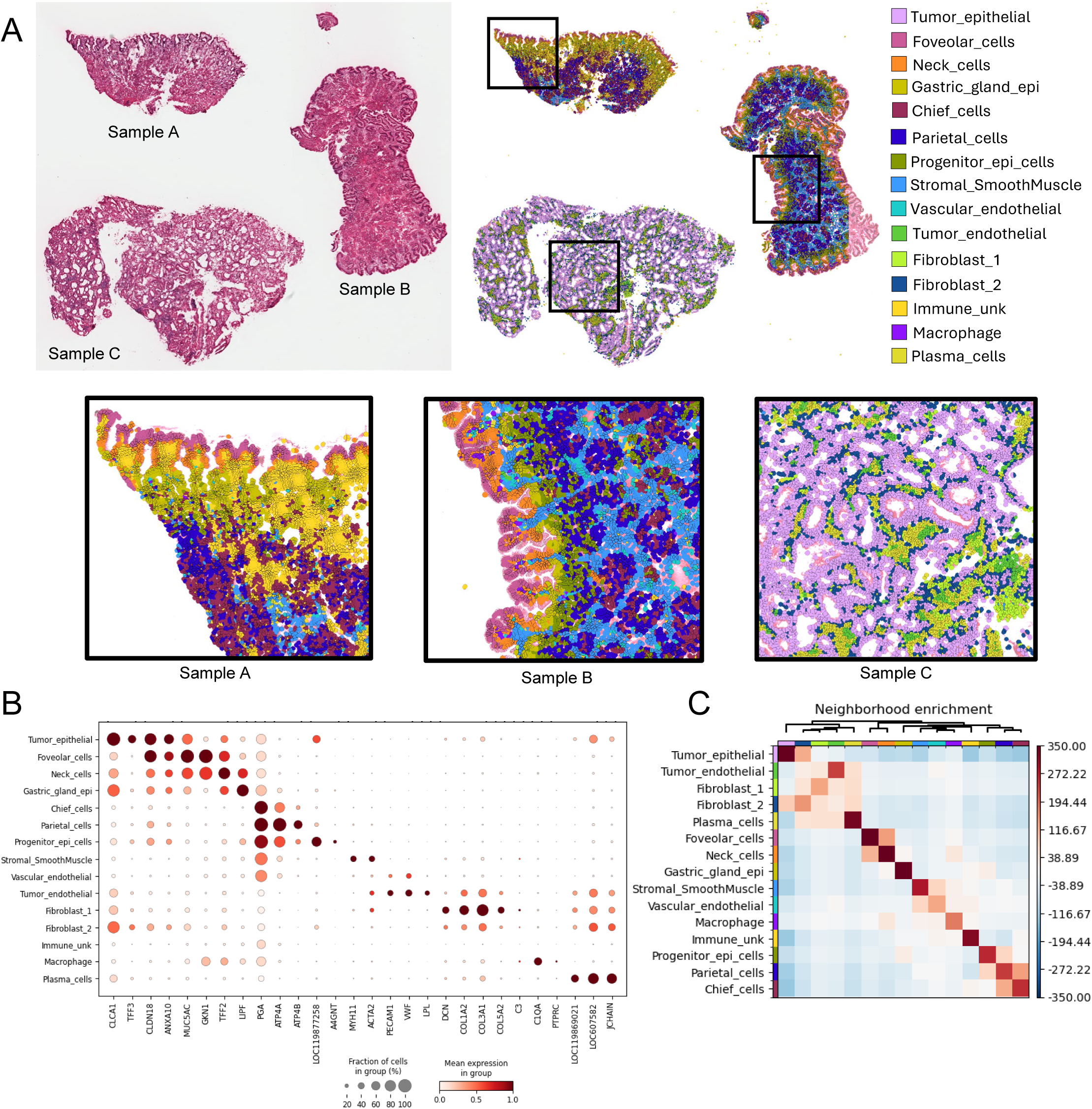
Spatial mapping of one gastric tumor and two normal stomach tissues. (A) Hematoxylin and eosin (left) stained tissue, paired with cell-segmented data (right), colored by unsupervised cluster. Samples A and B are examples of normal stomach tissue; Sample C displays neoplastic gastric tissue. Zoomed-in images corresponding to boxed regions above showing spatial localization of cell types. (B) Dotplot illustrating marker genes for each cell type. Human orthologs for LOCs are *MUC6* (*LOC119877258*), *IGHG1* (*LOC119869021*), *IGLC7* (*LOC607582*). (C) Neighborhood enrichment graph illustrating cell-cell proximity based on a connectivity graph of cell clusters. Colors indicate z-score.

Loss of parietal and chief cells observed in snRNA-seq data was supported at the tissue level by spatial transcriptomic analysis with tumor tissue exhibiting near-complete absence of marker gene expression for parietal (*ATP4A* and *ATP4B*) and chief (*PGA* and *PGB1*) cells. (Supplementary Figure 7).

### Bulk RNA-sequencing supports snRNA-seq findings and identifies Hippo pathway dysregulation

To validate snRNA-seq findings and expand the cohort size, we performed bulk RNA-seq on 10 gastric tumor and seven normal stomach tissue samples. PCA cleanly separated tumor and normal samples along the first principal component (Figure 7A). Differential gene expression analysis results from the bulk RNA-seq data were strongly correlated with the pseudobulk sn-RNAseq results across all genes (Pearson r = 0.715, 95% CI 0.708-0.721), with the correlation strengthening to r = 0.928 (95% CI 0.922-0.934) when restricted to genes significantly dysregulated in both datasets (n=2,267; Figure 7B).

**Figure 7.**
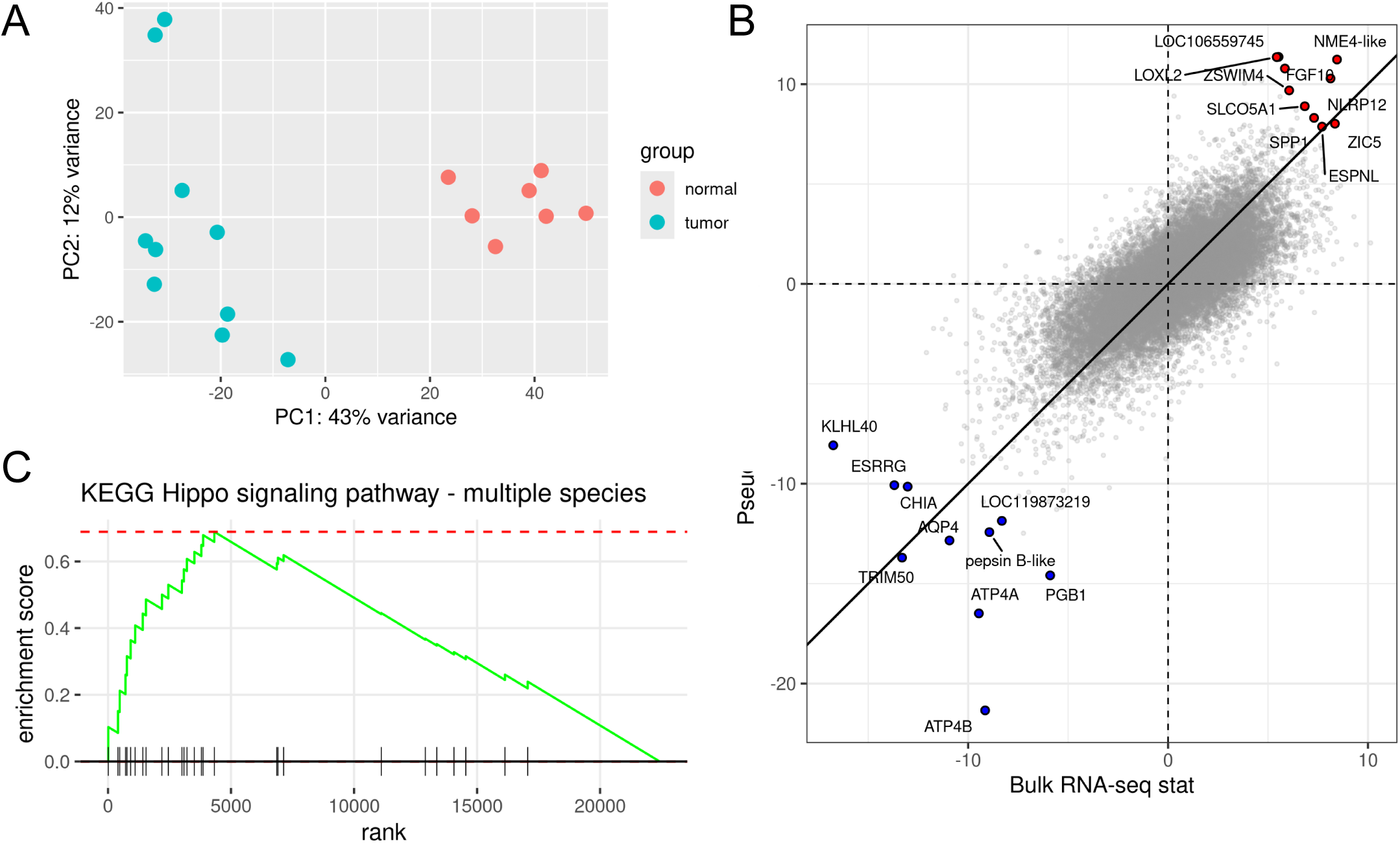
Bulk RNA-seq analysis from 10 tumor and 7 normal canine stomach samples. (A) PCA plot of the 500 most variable genes derived from the variance stabilizing method. Colors indicate stomach tissue phenotype. (B) Correlation plot of the Wald statistic reported by DESeq2 for bulk- and snRNA-seq tumor differential expression. Genes with the ten most positive and negative Wald statistics are labeled and colored by direction. (C) Gene set enrichment plot of the Hippo signaling pathway (KEGG ID: cfa04392) in the bulk RNA-seq data. Vertical black bars represent the rank of each gene in the pathway, the green line represents the running enrichment score, and the dashed line shows the maximum enrichment score.

Bulk RNA-seq differential expression recapitulated key pathways identified by snRNA-seq, including enrichment of the Hallmark epithelial-mesenchymal transition pathway (p_adj_ = 1.35×10^-27^) and depletion of oxidative phosphorylation (p_adj_ = 3.46×10^-36^) in tumor samples (Supplemental Tables 5-6). The bulk data additionally revealed significant enrichment of the Hippo pathway in tumors (KEGG cfa04392 p_adj_ = 4.27×10^-6^; Supplemental Table 7), with upregulation of the Hippo pathway transcriptional coactivators *YAP1* (p_adj_ = 5.28×10^-4^; log2FC = 1.40) and *WWTR1* (encoding TAZ; p_adj_ = 4.18×10^-5^; log2FC = 1.63), as well as their downstream target genes *CCN1* (*CYR61*; p_adj_ = 9.80×10^-6^; log2FC = 1.92) and *CCN2* (*CTGF*; p_adj_ = 4.23×10^-6^, log2FC = 2.38). The PI3K-Akt (p_adj_ = 4.01×10^-4^) and Wnt signaling (p_adj_ = 8.64×10^-4^) pathways were similarly enriched in tumor samples.

### Canine gastric cancer susceptibility genes display altered cell-type specific expression patterns

In the snRNA-seq data, we examined the cell-type-specific expression of canine gastric cancer susceptibility genes previously identified through our GWAS and genomic selection scan analyses in Belgian Tervuren and Belgian Sheepdogs^8^. Two genes identified through GWAS that had not been previously associated with gastric cancer, *PDZRN3* and *KLHL29*, were overexpressed in specific cell populations (Figure 8D). *PDZRN3*, the most significant GWAS association, was upregulated (p_adj_ = 3.00×10^-3^; log2FC = 1.81) in tumor endothelial cells displaying angiogenic transcriptional features (Figure 8A-C). *KLHL29* was significantly upregulated in tumor-derived neural, immune, and fibroblast populations, with the strongest upregulation observed in epithelial cells (p_adj_=1.50×10^-33^; log2FC = 4.98). *PLAU*, identified through our previous genomic selection scan analyses^8^, was upregulated across epithelial, endothelial, and immune cell compartments, with the strongest expression increase in epithelial cells (p_adj_ = 8.42×10^-10^; log2FC = 4.61; Figure 8E). Together, these findings link germline susceptibility loci to specific cellular contexts of dysregulation in canine gastric tumors and may inform mechanistic hypotheses for their roles in disease pathogenesis.

**Figure 8.**
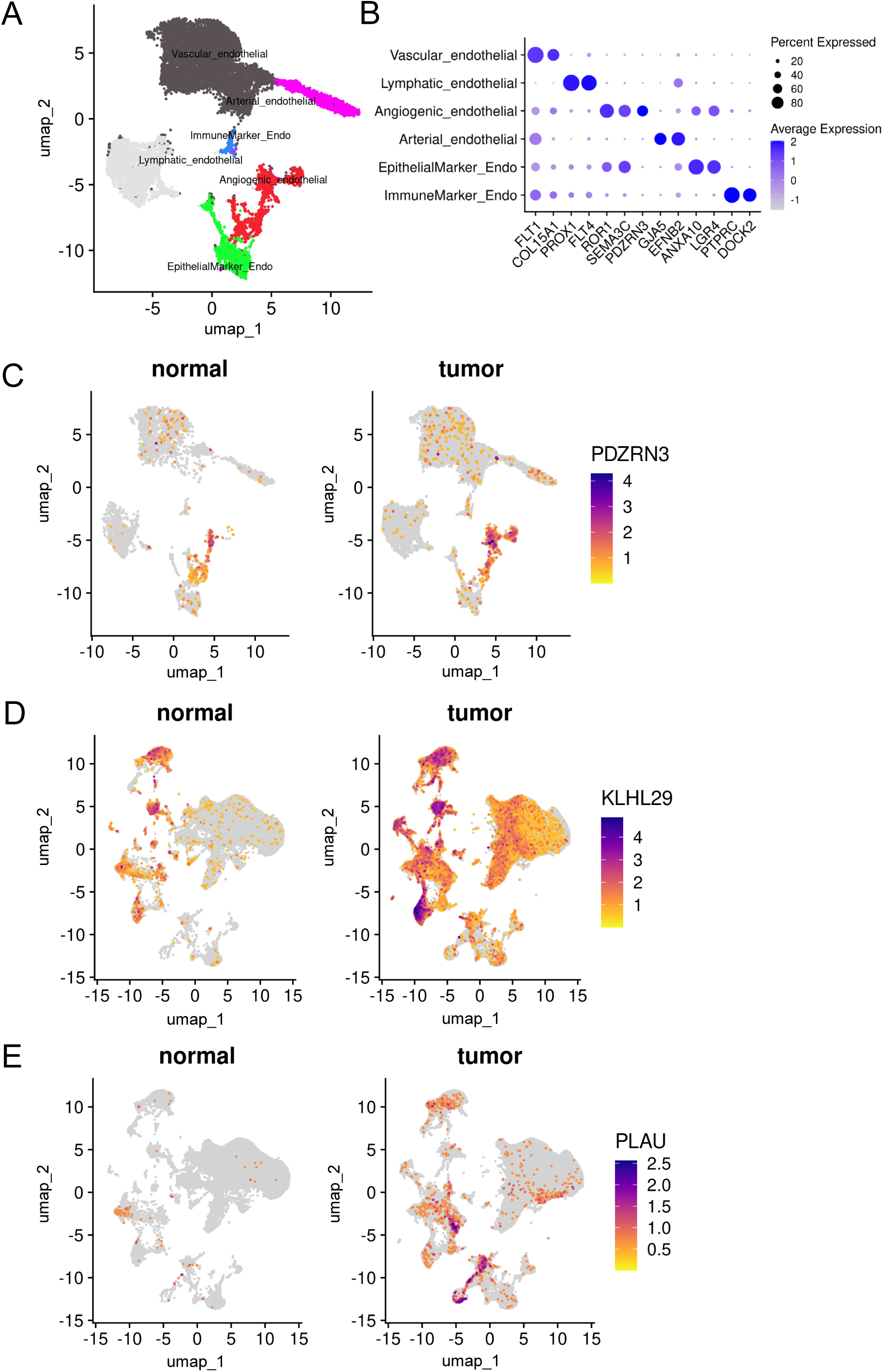
Characterization of endothelial cell populations and differential expression of germline susceptibility genes. (A) UMAP of subclustered endothelial cells (n = 14,808). (B) Dot plot showing expression and specificity of endothelial cell type marker genes. (C) UMAP of endothelial cells showing expression of *PDZRN3* across normal and tumor samples. (D) UMAP of all cell types illustrating the expression level of *KLHL29* and (E) *PLAU* in normal and tumor samples.

## DISCUSSION

We present the first transcriptomic characterization of canine gastric cancer, encompassing snRNAseq, bulkRNAseq, and single-cell resolution spatial transcriptomics of treatment-naive tumor and normal stomach tissues from Belgian Tervuren and Belgian Sheepdogs. Together, these data establish a high-resolution molecular atlas of canine gastric cancer, demonstrate substantial transcriptomic homology with human gastric cancer, and support the dog as a translationally relevant model for studying gastric cancer biology and evaluating new therapeutic strategies.

Canine tumors were characterized by significant loss of parietal and gastric chief cells, hallmarks of human gastric cancer, that normally function to secrete gastric acid and pepsinogen, respectively^23–25^. The depletion of these lineages in canine tumors, identified through snRNA-seq and supported by spatial transcriptomics, parallels gastric epithelial dedifferentiation observed in human gastric cancer. In humans, detection of anti-parietal cell antibodies and serum pepsinogen have demonstrated potential as biomarkers for early diagnosis^26–28^, and warrant evaluation in canine patients.

EMT was a dominant transcriptional program across multiple cell populations in canine gastric tumors. Under EMT, epithelial cells acquire mesenchymal features that promote motility and invasion, and EMT activity is associated with poor prognosis and metastasis in human gastric cancer^29–31^. We identified an EMT-like epithelial population expressing EMT transcription factors, as has been described in human gastric cancer single-cell studies^9^. Cell-cell communication analysis revealed substantially enhanced laminin and collagen signaling between epithelial cells and *POSTN+* matrix CAFs in tumors. *C3*+ CAFs were enriched in tumor samples and have been reported to drive EMT and cancer cell migration in human gastric cancer^16^. EMT signatures from tumor-derived fibroblast, epithelial, and immune cells, combined with epithelial-fibroblast crosstalk within laminin and collagen inferred signaling pathways, supports a stroma-driven EMT model, paralleling current understanding of EMT in human gastric cancer^30,32,33^.

Canine gastric tumors exhibited disruption in several core oncogenic pathways (Hippo, PI3K-Akt, Wnt) that are implicated in human gastric tumors driven by *H. pylori* infection, a major environmental risk factor associated with approximately 70-75% of human cases worldwide^34,35^. Hippo pathway disruption is a proposed primary mechanism by which *H. pylori* drives carcinogenesis in humans^36^. Bulk RNA-sequencing identified significant enrichment of the Hippo signaling pathway in tumors with upregulation of the transcriptional coactivators *YAP1* and *WWTR1* (TAZ) and their downstream targets *CCN1* and *CCN2*. PI3K-Akt and Wnt signaling, also activated by *H. pylori* in humans^36–38^, were similarly dysregulated in canine tumors. While *Helicobacter* species have not been associated with GC in dogs^4,39^, the disruption of these pathways across species suggests convergent oncogenic programs in gastric carcinogenesis, regardless of the inciting pathogen or environmental trigger. Further investigation of this mechanism will inform our understanding of human gastric cancer initiation.

In canine gastric tumor epithelium, we observed significant overexpression of *NLRP12*, *CDH6* and *CDH11*, all of which are upregulated in human gastric cancer tumors. *NLRP12* was the most significantly upregulated gene and correlates with poor prognosis in humans^40^. *CDH6* expression was nearly absent in normal samples but strongly expressed in tumor-epithelial cells. *CDH6* acts as an oncogene in human gastric cancer, where its expression distinguishes tumor from normal tissues with 90% specificity in TCGA data^42^, suggesting utility as a diagnostic biomarker. *CDH11*, encoding OB-cadherin, was also significantly upregulated in canine tumor samples and was predominantly expressed in the EMT-like epithelial cells, consistent with its established role in promoting EMT and gastric cancer progression in humans^43,44^. *CDH11* has been reported to facilitate interactions between gastric cancer cells and fibroblasts, leading to activation of YAP signaling^45^. Concurrent overexpression of *CDH6*, *CDH11*, and *CD44* has been described in human oral squamous cell carcinoma^46^.

*SPP1* was among the most significantly upregulated genes in immune cells from tumor samples and is commonly expressed by tumor-associated macrophages (TAMs) across cancer types^47–51^. Single-cell RNAseq studies in human gastric cancer have shown interaction of SPP1+ TAMs with CAFs^52^ and CD8+ exhausted T cells^47^. *SPP1* expression is increased in plasma, serum, and tumor tissue in humans with gastric cancer, and may have diagnostic value^53^. *CD109* was also overexpressed among the broad immune cell cluster and has been associated with EMT and an immunosuppressive TME in canine cancer, in addition to SPP1^54^. *OSMR*, which encodes the receptor for oncostatin M, was similarly upregulated. OSM signaling has been described as a key mediator of myeloid-stromal communication in human gastric cancer^55,56^.

Upregulation of these inflammation and immunosuppression associated genes together with the strong enrichment of TNFα/NF-κB and inflammatory response pathways supports the model of tumor-promoting inflammation as a fundamental feature of gastric cancer that is conserved between canines and humans. We also identified a specialized immune cell population of conventional dendritic cells (cDC1), expressing *XCR1*, which is an emerging target for tumor vaccine development^57–61^.

Spatial transcriptomics provided single-cell resolution of canine gastric architecture. In healthy tissue, spatial organization of epithelial populations recapitulated the canonical pit-to-base differentiation gradient of gastric fundic glands, paralleling human gastric mucosa^12^. Notably, we observed spatial co-localization of plasma cells, fibroblasts, and endothelial cells in tumor tissue, suggesting perivascular immune-stromal niches that warrant further investigation.

Expression of novel gastric cancer risk genes identified in our previous genomic analyses^8^ was increased in canine tumor samples. *KLHL29* was significantly overexpressed in multiple cell types, particularly epithelial cells. *KLHL29* is a member of the kelch-like gene family and a substrate adaptor of the Cullin3-RING ligases, which regulate degradation of oncoproteins and tumor suppressors in cancer^62^. While downregulation of *KLHL29* has been described in human breast cancer^63^, its role in gastric cancer has not been previously characterized, and the observed tumor-epithelial upregulation warrants further mechanistic investigation. *PDZRN3*, a ubiquitin E3 ligase and the only gene present within the most significantly associated GWAS locus, was overexpressed in angiogenic endothelial cell populations in canine tumors. In mice, *PDZRN3* overexpression in endothelial cells has been reported to disrupt intercellular junctions and increase vascular permeability^64^, suggesting a potential role in tumor vasculature. *PLAU*, identified through our previous genomic selection scan analysis^8^, was significantly upregulated in several cell types in canine tumors and is also overexpressed in human gastric cancer^65^. This work links germline susceptibility genes to the cell types in which they are aberrantly expressed and may provide insight into their potential functions in gastric cancer biology.

Limitations of this study include that while we identified over 40 distinct cell populations including rare states, the snRNA-seq and spatial transcriptomic cohorts were of modest size, and further studies with larger cohorts are required for robust validation. Future work should also include analyses stratified by tumor histological and/or molecular subtypes. While our cohort focused on Belgian Tervuren and Belgian Sheepdogs given the established breed predisposition to gastric cancer, future studies including additional breeds will be important for evaluating the generalizability of our findings.

We present the first single-nucleus, bulk, and spatial transcriptomic atlas of canine gastric cancer, providing high-resolution molecular characterization of this disease. Our findings nominate candidates for future diagnostic and therapeutic investigation, providing a foundation for veterinary clinical advances in canine gastric cancer. The conservation of cell type-specific transcriptional programs, oncogenic signaling pathways, and tumor microenvironmental features between canine and human gastric cancer supports the dog as a translationally relevant comparative oncology model.

## MATERIALS & METHODS

### Sample preparation for snRNA-seq

Nuclei were extracted from seven tumor and four normal stomach tissues, collected postmortem or upon diagnostic endoscopy in accordance with institutional protocols, and preserved in RNAlater solution with long-term storage at -80°C, excepting shipment. Tissues were dissociated on ice in Dounce tissue homogenizers using a digestion buffer (10mM Tris, pH 7.5; 20mM MgCl2; 142mM NaCl; 1mM CaCl2; 0.1% Tween 20; 1mM DTT; 0.3mg/mL Collagenase I; 1.5mg/mL Collagenase IV; 1X Halt Protease Inhibitor Cocktail; 0.2U/μL Protector RNase Inhibitor). Lysis buffer (10mM Tris, pH 7.5; 3mM MgCl2; 10mM NaCl; 0.1% Tween 20; 1mM DTT; 0.075% NP40; 0.2U/μL Protector RNase Inhibitor) was added to the homogenate, gently pipetted 15 times, then incubated on ice for 5 minutes. The lysate was then transferred to a 2mL Lo-Bind tube and centrifuged at 300 × g for 10 minutes at 4°C before being resuspended in wash buffer (1X PBS; 0.75% BSA; 1mM DTT; 0.1% Tween 20; 0.2U/μL Protector RNase Inhibitor) filtered through a 35μm mesh. Nuclei were subjected to two more rounds of centrifugation (300 × g for 10 minutes at 4°C) and resuspension in wash buffer before a final centrifugation (300 × g for 10 minutes at 4°C), resuspension in resuspension buffer (1X PBS; 0.75% BSA; 0.2U/μL Protector RNase Inhibitor, and filtration through a 35μm mesh. Single cell suspensions were run on a Chromium X instrument, targeting 10,000 nuclei per sample. Libraries were prepared with the Chromium GEM-X Single Cell 3′ v4 Assay, following the 10x Genomics standard protocol (user guide CG000731, RevB) using 12-13 cycles of cDNA amplification. Sample quality was assessed using a Qubit (DNA HS kit; Thermo Fisher) to determine concentration and a Fragment Analyzer (Agilent) to confirm fragment size integrity. Libraries were sequenced (2×150bp) on a NovaSeqX. Libraries and sequencing were generated by the Cornell BRC Genomics Facility (RRID:SCR_021727).

### Sample preparation for bulk RNA-seq

Bulk RNA-seq was performed for 10 treatment-naive tumor and seven normal stomach samples collected postmortem, preserved in RNAlater, and stored at -20 or -80^°^C. Total RNA was extracted using Trizol (Thermo Fisher Scientific, Waltham, MA). Approximately 200 mg each tissue was minced, added to 1 ml Trizol (Thermo Fisher Scientific, Waltham, MA) in a 1.5 ml tube and homogenized with a disposable pestle. After centrifugation for 10 minutes (min), 12,000 x g at 4^°^C, lysates were transferred to clean 1.5 tubes, 200 µl chloroform isoamyl alcohol (Thermo Fisher Scientific) were added and tubes were shaken by hand for 30 seconds. Tubes were centrifuged for 15 min, 16,000 x g at 4^°^C, aqueous layers were moved to clean 1.5 ml tubes and equal volumes of 100% EtOH (Decon Labs, King of Prussia, PA) were added. Sample material was transferred to Zymo-Spin IIICG columns from a Zymo Quick-RNA Miniprep Plus Kit (Zymo Research, Irvine, CA). DNase1 treatment, sample washing, and RNA elution were carried out following manufacturer’s instructions. RNA concentrations were determined on a Qubit fluorometer (Thermo Fisher), and quality was assessed using a 5200 Fragment Analyzer system (Agilent Technologies, Santa Clara, CA). NEBNext Ultra II [Directional] RNA stranded polyA enrichment libraries (2×150 paired-end) were prepared by the BRC Genomics core and sequenced on an Illumina NovaSeqX.

### Visium HD 3′ spatial transcriptomics

Tissues were collected as part of standard diagnostic endoscopy while animals were under anesthesia or during postmortem autopsy. Procedures were performed by a licensed veterinary clinician (PJJM). The complete stomach was examined macroscopically, and biopsies were taken from the edges and center of visually abnormal tissue, or a normal region distant from any abnormal tissue. Samples were embedded in Optimal Cutting Temperature (OCT) Compound using an isopentane bath cooled with liquid nitrogen immediately following collection and stored at -80°C, excepting shipment on dry ice to Cornell University.

Three gastric tissues were utilized for spatial transcriptomics (Figure 6). All three samples were derived from dogs diagnosed with gastric cancer. Sample A was collected from a tumor-free region of the stomach and was histologically normal. Sample B, sectioned on a slightly oblique plane, exhibited no evidence of neoplasm. Sample C clearly exhibited disorganization characteristic of neoplastic tissue.

Spatial transcriptomics data were collected using the Visium HD 3’ Gene expression platform. Fresh-frozen, OCT-embedded tissues were sectioned at –20°C using a Microm HM505E cryostat with a thickness of 10μm. All three tissues were sequentially sectioned, mounted on the glass slide, and warmed, with the microtome blade repositioned to a fresh cutting edge upon tissue exchange. The slide was then processed following the Visium HD 3’ Fresh Frozen Tissue Preparation Handbook (CG000804, Rev B) and Visium HD 3’ Spatial Gene Expression User Guide (CG000805, Rev B) and sequenced on the NovaSeqX platform.

### Data processing for snRNA-seq

CellRanger v.9.0.1^66^ was used to align raw fastqs to the canine reference genome (UU_Cfam_GSD_1.0_ROSY) with mitochondrial sequence and annotation derived from CanFam6. Transcript counts (nCount_RNA), number of detected genes (nFeature_RNA), mitochondrial gene fraction, and ribosomal gene fraction were calculated for each nucleus. Nuclei from each sample were excluded if they contained >10% mitochondrial fraction and were further filtered using data-driven quality control (DDQC)^67^, which performs unsupervised clustering and removes nuclei with nCount_RNA, nFeature_RNA, mitochondrial gene fraction, and/or ribosomal gene fraction more than two absolute standard deviations away from the median. DoubletFinder^68^ was used to identify putative doublets, which were removed.

Gene counts were normalized with sctransform^69^, the datasets were integrated with Seurat^70^ using the reciprocal principal component analysis (RPCA) method, and the nuclei were clustered with FindClusters. Dimensionality reduction was performed using the top 25 PCs and visualized via UMAP.

Using a clustering resolution of 0.4, differential gene expression analysis with FindAllMarkers was performed using only.pos=TRUE and a log_2_ fold change of 0.2 to enable cell type assignment based on cluster-enriched markers. The top 20 marker genes per cluster were examined to determine cell type. Additionally, cell types were validated using canonical marker genes (e.g., endothelial: *VWF*, *PECAM1*; immune cells: *PTPRC*; fibroblasts: *ACTA2*, epithelial: *CLDN18*). Further cell type classification was performed by subsetting each broad cluster and reclustering by performing normalization, integration, and clustering of the subset nuclei. Again, cell type assignment was based on the top 20 cluster-enriched markers.

Cell type specific differential gene expression analysis was performed using Seurat’s AggregateExpression to pseudobulk the data and subsequently analyzed in DESeq2^71^.

### Cell type proportion analysis of snRNA-seq data

Cell type proportion differences were calculated using sccomp^72^. The cell type annotated Seurat object was provided directly to sccomp, and outliers were removed with sccomp_remove_outliers. The proportion of each subcluster was calculated compared to its broad cell annotation cluster (*i.e.*, cancer-associated fibroblast proportion out of all fibroblast cells). Differences in cell type proportion between tumor and normal samples were considered significant if FDR ≤ 0.05.

### Cell Chat intercellular communication analysis

Cell-cell communication was assessed using the R package CellChat v2.2.0^73^ using CellChatDB human. Tumor and normal data were analyzed separately using normalized gene expression data and predefined cell-type annotations prior to integration with mergeCellChat(). Effect of cell proportion in each cell group was incorporated using population.size = TRUE. All other parameters were default, as recommended in the CellChat documentation.

### Data processing for bulk RNA-seq

On average we achieved 72M paired reads per sample. Raw paired-end fastq files were processed using nf-core/rnaseq 3.19.0^74^. In short, adapter sequences and low-quality bases were removed with Trim Galore!^75^, reads were aligned to the UU_Cfam_GSD_1.0_ROSY reference genome using the NCBI annotation (release 106) with STAR^76^, and gene and transcript counts were calculated with Salmon^77^.

### Differential gene expression analysis of bulk RNA-seq data

Transcript-level counts were imported into Rstudio with tximport^78^ and converted to a DESeqDataSet using tumor vs. normal condition as a covariate. Genes with fewer than 10 counts across all samples were excluded. Samples were visualized in a PCA using the regularized log transformation to identify any outliers. Differential expression analysis was performed with DESeq2, and genes with an adjusted p-value < 0.05 and │log2FC│ ≥ 1 were considered differentially expressed. A clustered heatmap using the expression of the top 500 genes was created using pheatmap with the ward.D2 clustering method. Pearson correlations between the bulk- and snRNAseq datasets were calculated using the Wald statistic reported by DESeq2, which represents gene expression normalized by its standard error.

### Gene set enrichment analyses

Gene set enrichment analysis was performed on the bulk RNAseq and pseudobulked snRNA-seq broad cell populations. All data were ranked using the Wald statistic calculated by DESeq2. Enrichment of KEGG pathways was analyzed in clusterProfiler^79^ using the *Canis lupus familiaris* database. The eps option was set to 0 to allow for precise p-value calculation.

For analysis of enriched Hallmark pathways from MSigDB^80^, gene ortholog data were downloaded from NCBI (https://ftp.ncbi.nlm.nih.gov/gene/DATA/ gene_orthologs.gz; accessed May 31, 2026), and canine and human orthologs were extracted. Canine gene names were updated to match the human ortholog (Supplemental Table 8) for gene set enrichment analysis and gene set co-regulation analysis via Bioconductor’s fgsea package^81^. The January 2026 release of the human gene sets was used for all analyses.

### Data processing for spatial transcriptomics

Fastq files were aligned to UU_Cfam_GSD_1.0_ROSY, with CanFam6 mitochondrial sequence and annotation using 10X Genomics Space Ranger v4.0.1 count pipeline using default parameters. Cell-segmented data were analyzed using Loupe Browser v9.0.0. Graph-based clusters were annotated by integrating spatial localization of canonical marker gene expression with cluster-specific differential expression profiles.

For downstream analysis in python, 2μm binned data were mapped to segmented cells using the barcode_mappings.parquet file, output by Space Ranger. Neighborhood enrichment analysis was performed using Squidpy 1.6.1^82^ with 1000 permutations and default parameters. Ward variance minimization algorithm was used to generate a hierarchical clustering map for plotting with squidpy.pl.nhood_enrichment.

## DATA AVAILABILITY

Raw sequencing data generated herein will be made available in the NCBI Sequence Read Archive upon publication under accession number PRJNA1480067.

## ACKNOWLEDGEMENTS

We are grateful to the owners and veterinarians who provided samples and clinical data for this study. We thank Bo Shui for his expertise in designing the nuclei isolation protocol and Yingshan Fang for her contributions to nuclei isolation and preliminary tissue quality control. We thank Dr. Chris Champion for histological review of the spatial transcriptomic data. Sequencing services were provided by the Cornell University Biotechnology Resource Center Genomics Core facility (RRID:SCR_021727), with snRNAseq and Visium HD 3′ library preparation performed by Dr. Yassi Hafezi, and by the Transcriptional Regulation and Expression Core Facility (RRID:SCR_022532), which performed bulk RNA-sequencing. This work was supported in part by the Cornell Center for Vertebrate Genomics Seed Grant, Albert C. Bostwick Foundation, and Belgian Sheepdog Club of America and individual donations.

