## Supplemental Figures for "A multi-modal transcriptomic atlas reveals the cellular and spatial landscape of canine gastric cancer"

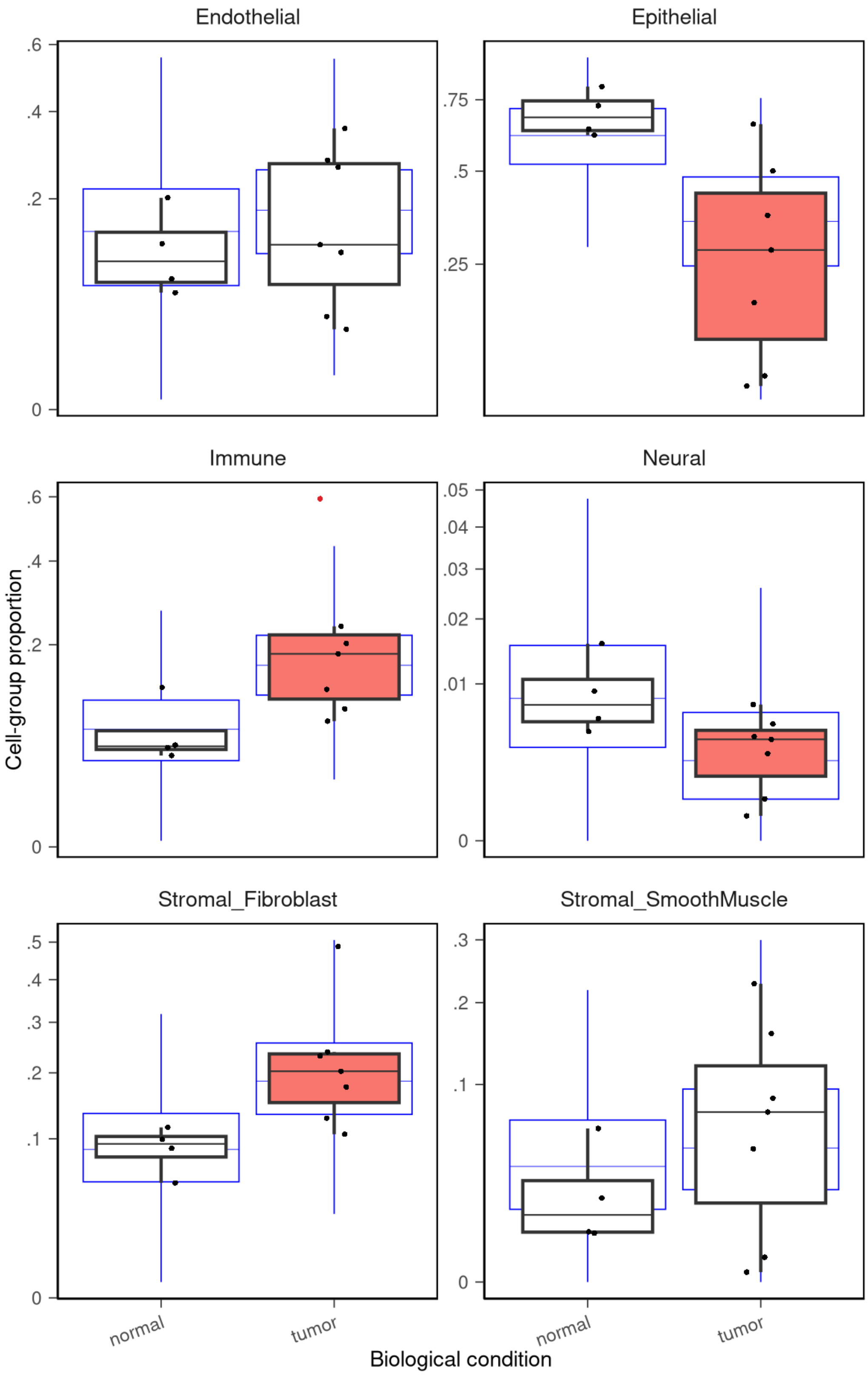


**Supplemental Figure 1. Differential cell-type composition analysis of the broad annotated cell populations using sccomp.** Black boxplots represent the observed data, and blue boxplots represent posterior predictive distributions. Each sample (4 normal, 7 tumor) is plotted (dot), with red dots indicating outliers that were removed prior to statistical testing. Boxplots in pink are significantly different from the comparison group at FDR < 0.05.

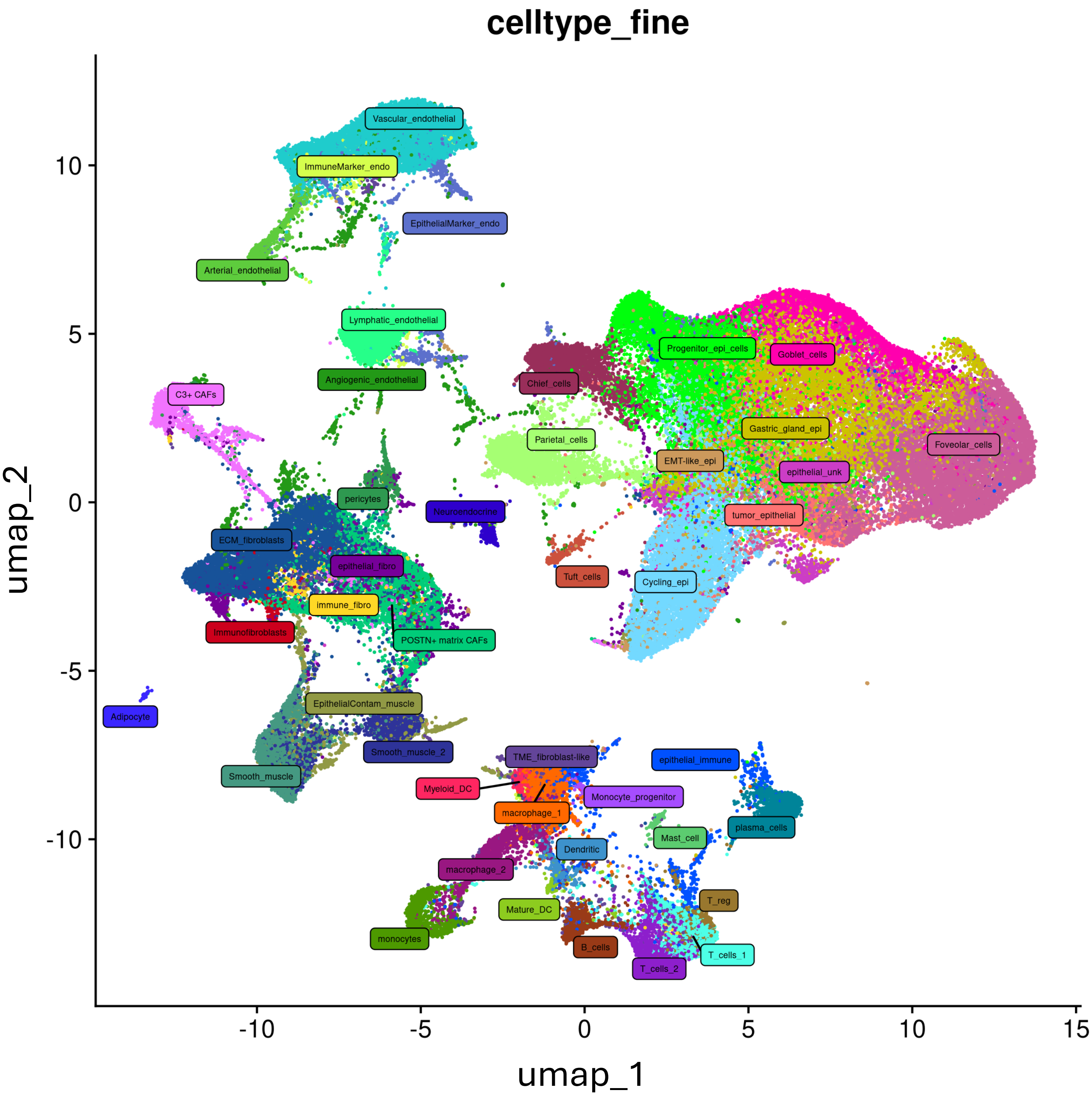


**Supplementary Figure 2. UMAP annotated with all subclustered cell types.** Cell annotations resulting from independent subclustering of each of the 7 broad cell categories are mapped back onto the overall UMAP (107,085 nuclei).

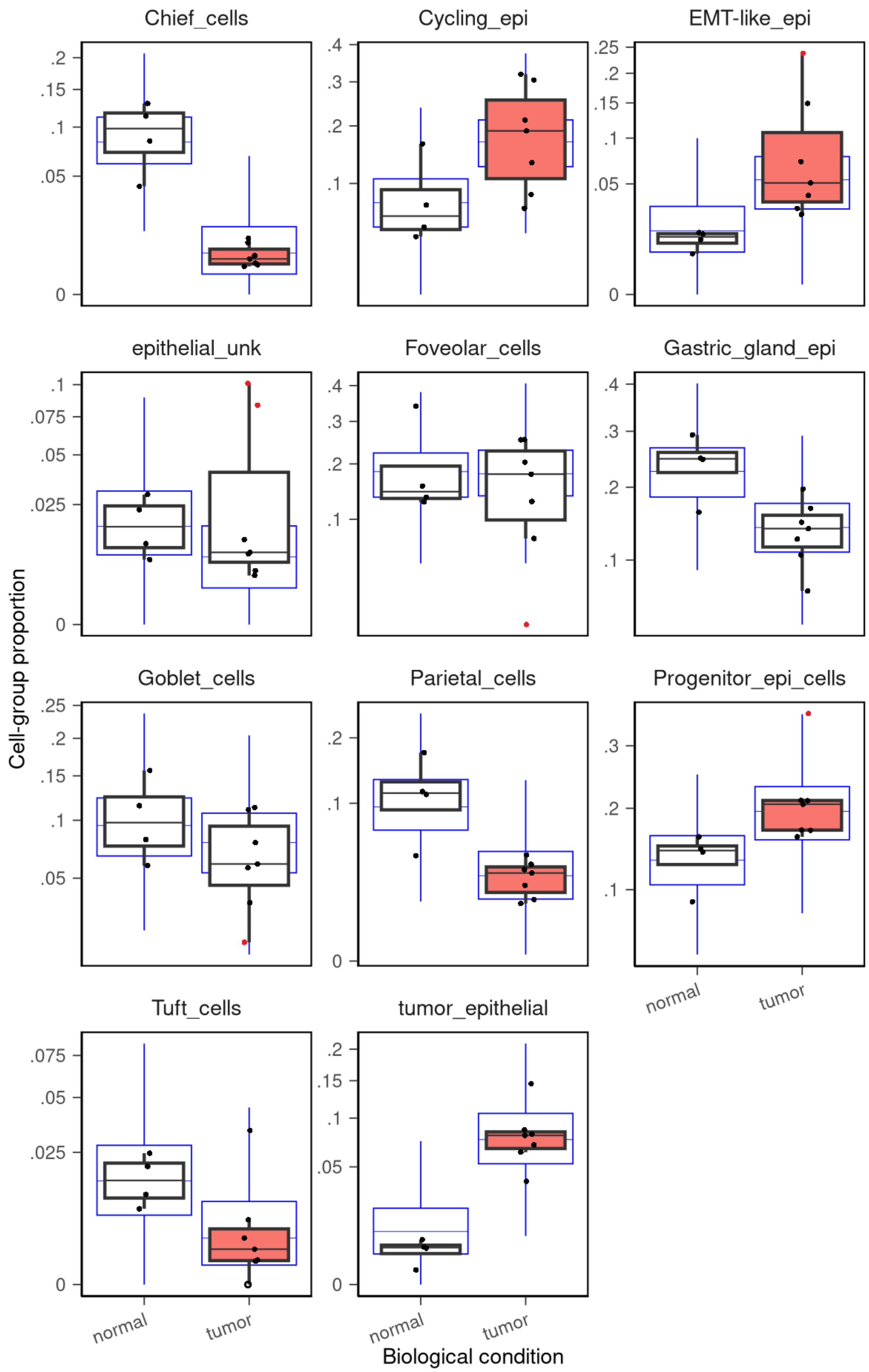


**Supplemental Figure 3. Differential cell-type composition analysis of the epithelial subclusters using sccomp.** Black boxplots represent the observed data, and blue boxplots represent posterior predictive distributions. Each sample (4 normal, 7 tumor) is plotted (dot), with red dots indicating outliers that were removed prior to statistical testing. Boxplots in pink are significantly different from the comparison group at FDR < 0.05.

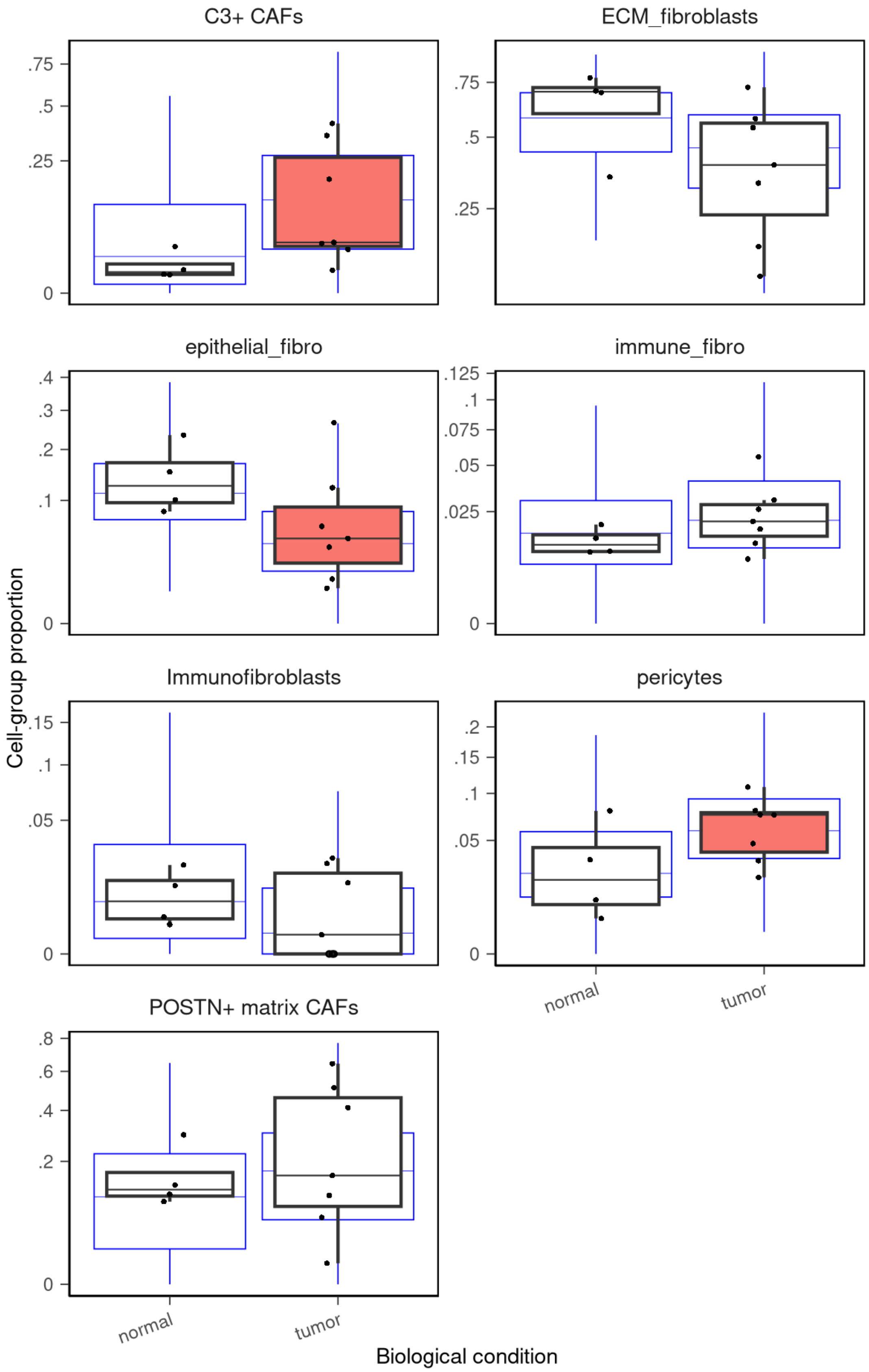


**Supplemental Figure 4. Differential cell-type composition analysis of the fibroblast subclusters using sccomp.** Black boxplots represent the observed data, and blue boxplots represent posterior predictive distributions. Each sample (4 normal, 7 tumor) is plotted (dot), with red dots indicating outliers that were removed prior to statistical testing. Boxplots in pink are significantly different from the comparison group at FDR < 0.05.

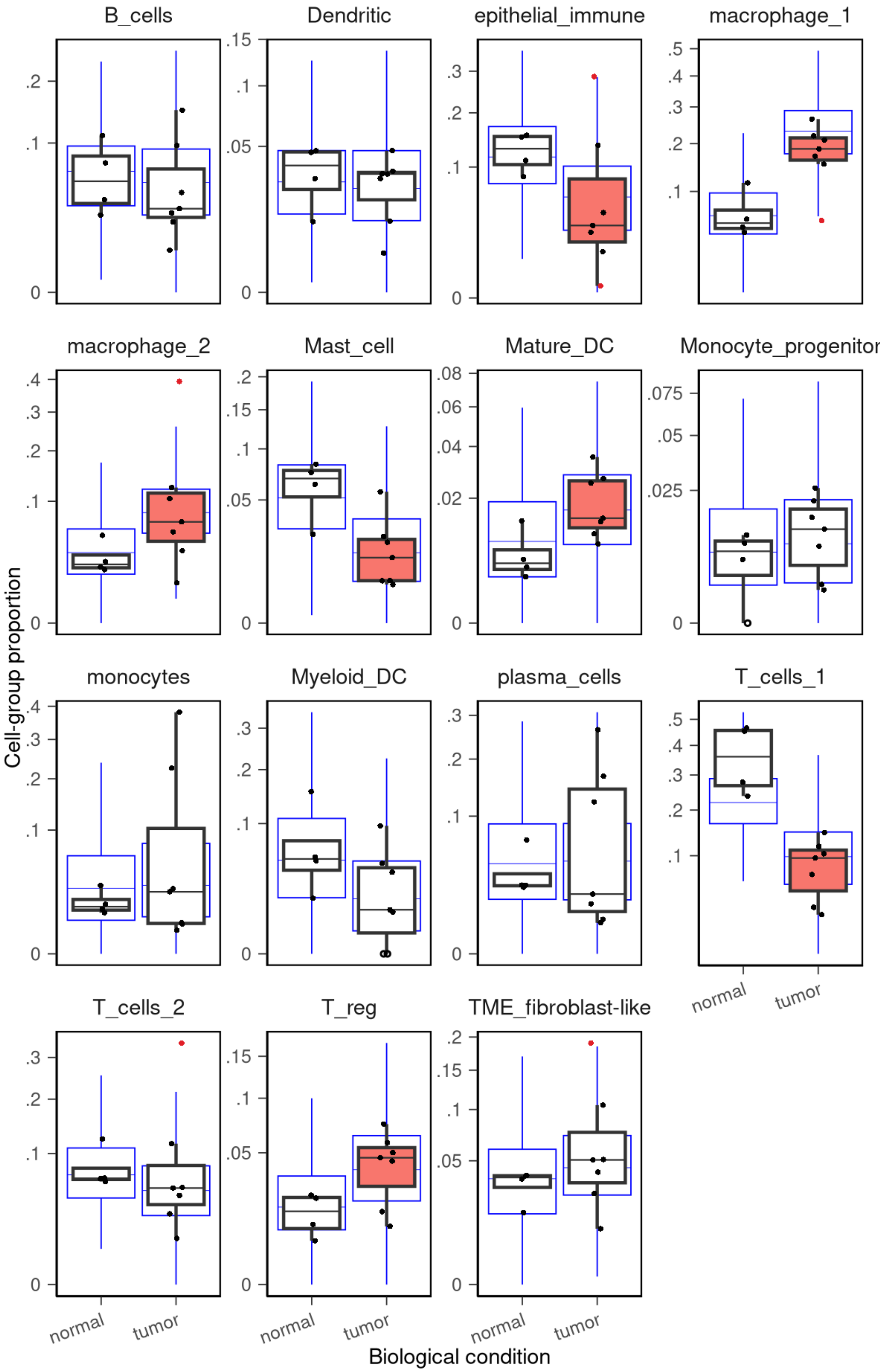


**Supplemental Figure 5. Differential cell-type composition analysis of the immune subclusters using sccomp.** Black boxplots represent the observed data, and blue boxplots represent posterior predictive distributions. Each sample (4 normal, 7 tumor) is plotted (dot), with red dots indicating outliers that were removed prior to statistical testing. Boxplots in pink are significantly different from the comparison group at FDR < 0.05.

A

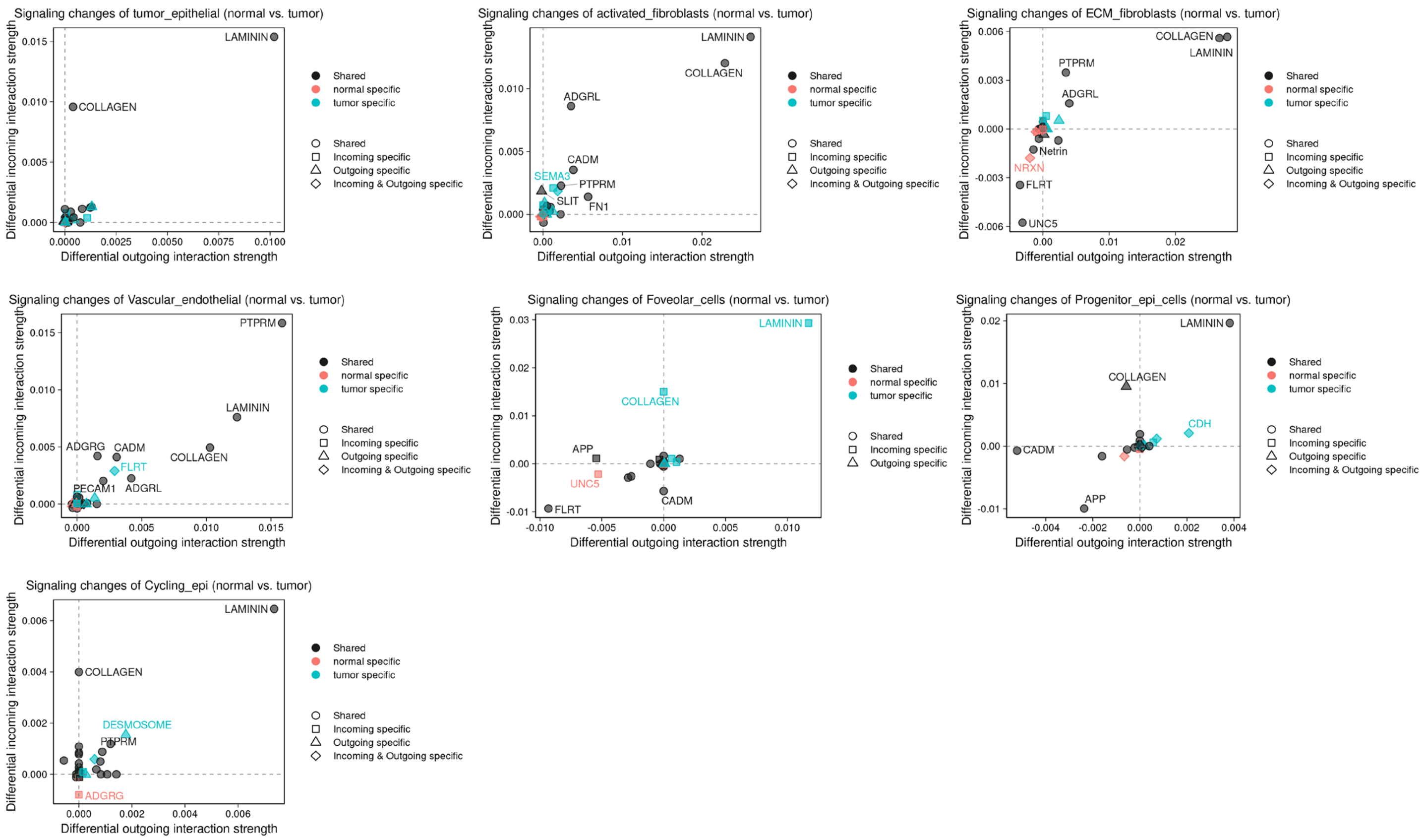


B

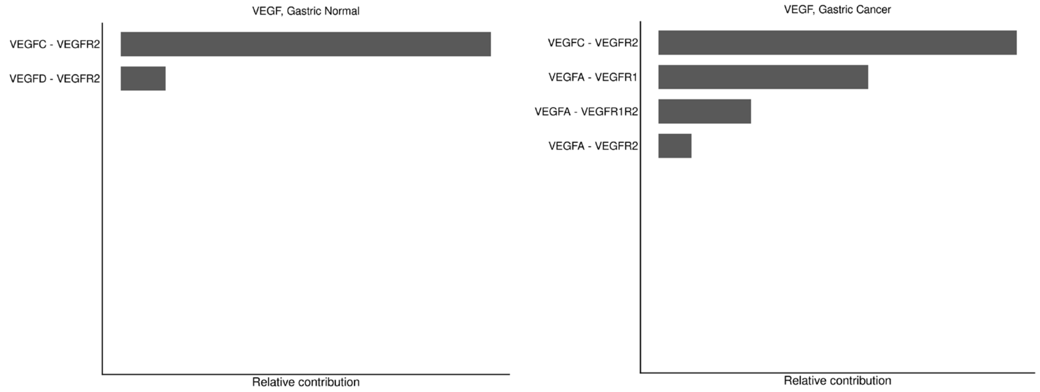


**Supplementary Figure 6. Differential outgoing and incoming signaling pathways within select cell populations.** (A) Scatter plots illustrate differential signaling patterns within tumor epithelial, POSTN+ matrix CAFs, ECM fibroblasts, vascular endothelial, foveolar, progenitor epithelial, and cycling epithelial cells. Positive values indicate an increase in signaling in tumor samples compared to normal. (B) Contribution of each ligand-receptor pair within the inferred VEGF signaling network for normal (left) and tumor (right).

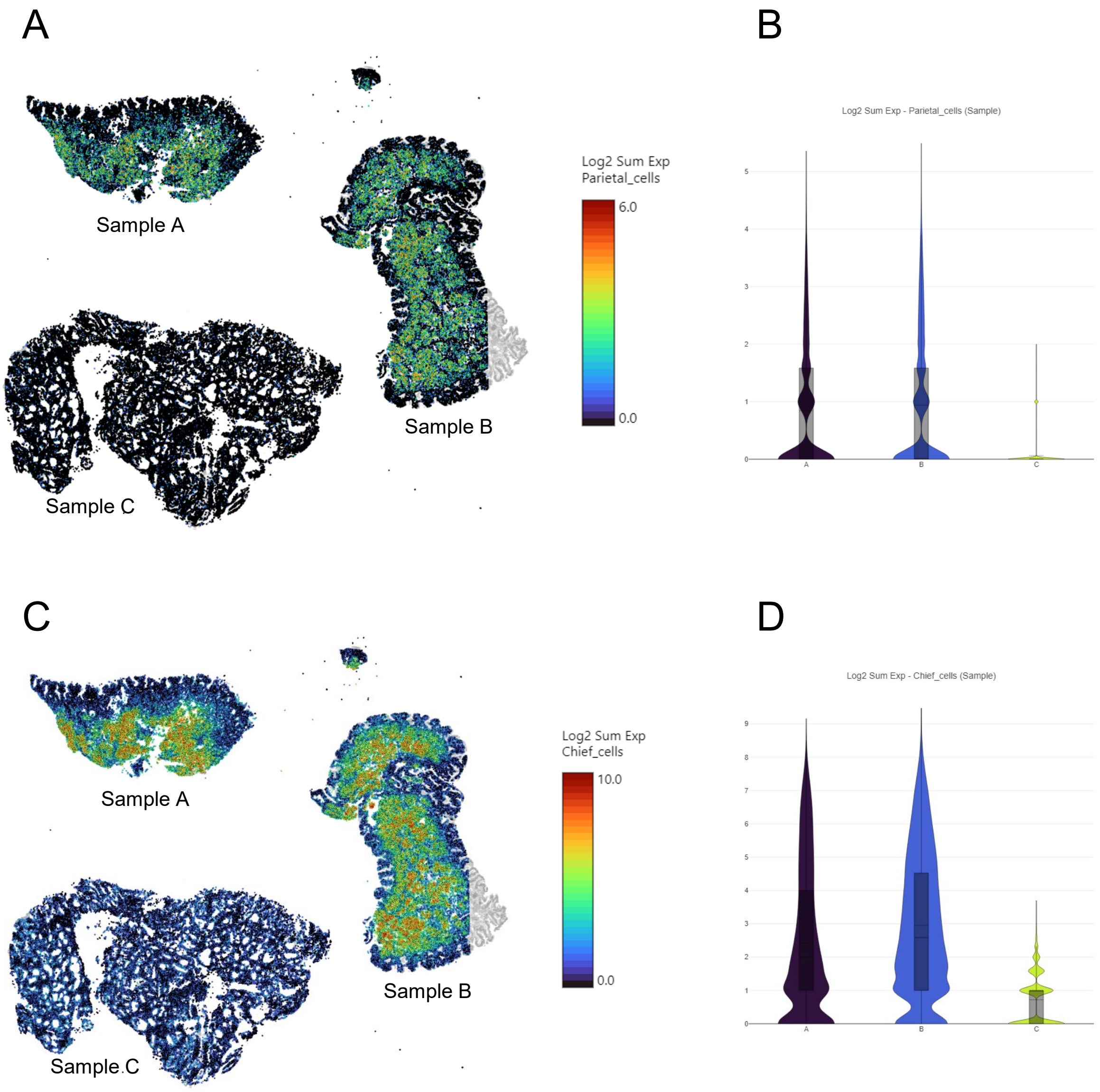


**Supplementary Figure 7. Loss of parietal and chief cell marker gene expression in gastric cancer.** (A) Spatial transcriptomic data illustrating summed expression of parietal cell (A-B) and chief cell (C-D) marker genes across normal (Samples A &B) and tumor (Sample C) tissues. Spots are colored by log2-transformed sum of expression across key marker genes for parietal (*ATP4A*, *ATP4B*) and chief (*PGA*, *PGB1*) cells.
